# Delineating the Spatial Patterns of Cell Type–Specific Stemness Using a Statistical Deconvolution Model

**DOI:** 10.1101/2024.12.29.630672

**Authors:** Eric J. Li, Liang Li, Ziyi Li

## Abstract

**Summary:** Cellular stemness is a cells ability to self-renew and differentiate into specialized cell types. In spatial transcriptomics (ST), researchers can use gene activity to quantify stemness and the associated spatial locations, allowing for the investigation of cellular profiles in their natural context. However, many ST platforms provide data with low resolution, and each data spot (i.e., spatial location) contains multiple cell types, which hinders the investigation of cell type–specific (CTS) behaviors, such as cell stemness, at different spatial locations. We developed a bivariate kernel-weighted regression method with constrained optimization to estimate CTS stemness as a function of spatial location and developed accompanying visualization tools. Through simulation studies and an application to real breast cancer ST data from the 10x Genomics Visium platform, we demonstrated that our method can accurately estimate CTS stemness and help shed light on the interplay among cell type, tissue structure, and stemness.

**Availability and implementation:** An R Shiny app implementing the proposed method is available at GitHub (https://github.com/ericli0480/stemness-deconvolution/).

**Supplementary information:** Available in a separate PDF file

## 1 Introduction

Cellular stemness represents a cell's ability to self-renew and differentiate into specialized mature cells. Totipotent stem cells and embryonic cells generally have high stemness, whereas muscle cells, nerve cells, and blood cells generally have low stemness. In contrast, most cancer cells exhibit a loss of differentiation (Friedmann-Morvinski and Verma 2014, Herreros-Villanueva et al. 2013). However, subpopulations of tumor cells called cancer stem cells often have self-renewal capabilities and are responsible for the long-term maintenance of tumor growth (Batlle and Clevers 2017). Moreover, cancer progression may be accompanied by cells' acquisition of differentiation potential, which can drive treatment resistance and disease relapse (Thorgeirsson 2016). In cancer research, analyzing cellular stemness can clarify the differentiation behavior of various cell types and show how the stemness varies according to the tissue location and other tissue microenvironmental factors. It may also provide insights into potential usage for differentiation therapy in cancer research.

To analyze cellular stemness, Gulati et al. (2020) developed Cellular (Cyto) Trajectory Reconstruction Analysis using gene Counts and Expression (CytoTRACE), a transcriptome-based computational algorithm that predicts differentiation states from single-cell RNA sequencing (scRNA-seq) data. However, scRNA-seq measures the gene expression of each cell without information about the cell's surrounding environment. The lack of spatial information prevents the investigation of a spatially heterogeneous tumor structure and its associated stemness pattern.

Recent advancements in spatial transcriptomics (ST) technology have enabled researchers to simultaneously measure transcriptomic gene expression and determine its spatial location. Some popular ST platforms, such as GeoMx (NanoString) and Visium (10x Genomics), measure the gene expression counts across a dense grid of spatial locations (“spots”) in a tissue section, thereby revealing the transcriptomic activity on a background of tissue features. However, these platforms have a low resolution of cells at each spot. For example, the Visium ST platform can capture a 6.5 mm × 6.5 mm tissue area that is covered with a two-dimensional array comprising 4,992 spots, each of which is 55 µm in diameter; the horizontal and vertical center-to-center distances between adjacent spots are all 100 µm. However, most cells are substantially smaller than 55 µm. Therefore, any spot-level gene expression data, including the cell stemness, reflects an average of approximately 10 to 30 cells in that spot. Although it is reasonable to assume that cells of the same type within a small neighborhood of tissue have the same stemness, the stemness of different types of cells within that neighborhood may differ. This potential variability makes it difficult to interpret spot-level data. We define “spot-level stemness” to be the average stemness of all the cells in each spot, the “cell-type specific stemness (CTS stemness)” to be the average stemness of all the cells of a specific cell type within each spot.

Statistical methods have been developed for the CTS estimation of the genetic features of scRNA-seq data (Rahmani et al. 2019, Jin et al. 2021) and bulk gene expression data (Feng et al. 2023). In these data, the cellular spatial location in the tissue is obscured; therefore, the spatial effect, if any, is treated as a random variation of the data. Using ST provides an opportunity to separate the spatial effect from other variation of the data, which allows the study of CTS features in different cell types and different spatial locations. To our knowledge, no existing statistical methods fully leverage the spatial correlation among spots and estimate the CTS stemness using ST data.

This paper presents a new statistical method estimating CTS stemness as a function of the spatial location. We also performed a simulation study to validate the performance of the proposed method and used the method to analyze Visium ST data from a breast cancer sample. We developed a web application that does not need additional programming to facilitate the use of this new method by biological laboratories. Although our method was developed to estimate CTS stemness, the framework is applicable to estimating any CTS genomic features, such as gene expression or pathway activities, with incorporation of spatial information.

## 2 Methods

### 2.1 Model Assumptions

Suppose the spot-level stemness and cell-type proportions were estimated using CytoTRACE and conditional autoregressive-based deconvolution (CARD) (Ma and Zhou 2022; Online Appendix) respectively. CytoTRACE quantifies stemness as a number between 0 and 1; higher numbers indicate higher stemness, i.e., a stronger potential for differentiation. In our model, we assume that the CTS stemness varies continuously across spots and that the stemness is more similar among adjacent spots and more different among distant spots. This smoothness assumption is well supported by biological evidence (Ståhl et al. 2016, Schürch et al. 2020) and is used in ST analyses (Ma and Zhou 2022). We also assume that the spot-level stemness is the weighted average of the CTS stemness, with the weights being the cell type proportions, at that spot. This admixture assumption is also commonly used in other CTS analyses (Rahmani et al. 2019, Jin et al. 2021, Feng et al. 2023). The two modeling assumptions above support the use of a kernel-weighted regression method to estimate the CTS stemness by spot (Wasserman 2005).

### 2.2 Statistical Model and Deconvolution Method

Let *N* spots of a tissue section be indexed by *i* = 1,2, … , *N*. Each spot contains many cells belonging to *K* cell types. We denote the cell-type proportions at the *i*^th^ spot by {*W*_*ik*_; *k* = 1,2, … · *K*}, where 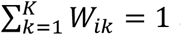 and all *W*_*ik*_ ≥ 0. The stemness at spot *i* is denoted by *Y*_*i*_, a number between 0 and 1. We denote the stemness of cell type *k* at spot *i* by *α*_*ik*_. Let *u*_*i*_ and *v*_*i*_ denote the horizontal and vertical coordinates of spot *i*, respectively; *α*_*ik*_ can also be written as a function of these coordinates: *α*_*ik*_ = *α*_*k*_(*u*_*i*_, *v*_*i*_). We estimate *α*_*k*_(*u, v*) as a smooth bivariate surface for each cell type. Our model assumes that

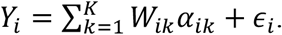

The residual *ϵ*_*i*_ is the mean-zero random noise that captures any deviation from the admixture assumption owing to variability in the cell-type proportion and spot-level stemness.

To estimate *α*_*k*_(*u, v*) for a given spatial location (*u, v*), we use a Gaussian kernel with bandwidth *h*. The Euclidean distance between spot *i* and the spot location (*u, v*) is

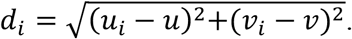

The kernel weight *ω* of spot *i* is

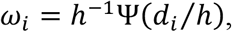

where ψ is the kernel function, taken here to be the density function of a standard Gaussian distribution. We simplify the notation by *α*_0*k*_ = *α*_*k*_(*u, v*). We can estimate {*α*_0*k*_; *k* = 1,2, … *K*} by minimizing the weighted least squares,

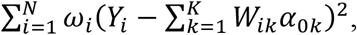

subject to the *K* inequality constraint that 0 ≤ *α*_0*k*_ ≤ 1. This constrained optimization problem can be solved using quadratic programming, which is available in the R package `quadprog`. We ran the procedure above *N* times, each time designating (*u, v*) to be a different spot on the tissue section. This algorithm produced *K* estimations of CTS stemness at each spot, denoted by 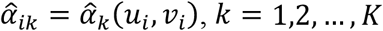.

### 2.2 Bandwidth Selection with Cross-validation

Selecting the bandwidth *h* is critical to kernel-weighted regression because of the bias–variance trade-off in nonparametric smoothing problems (Wasserman 2005). We selected *h* by using leave-one-out cross-validation. This iterative procedure had two steps in each iteration. First, we retained spot *i* for the validation data, and all other *N* − 1 spots for the training data. We used the training data to estimate the CTS stemness at (*u*_*i*_, *v*_*i*_), denoted by 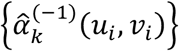. Second, we calculated the predictive accuracy on the validation spot *i* by 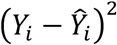 where 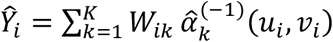. The procedure was repeated for all *N* spots, and the average prediction accuracy was defined as 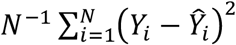. This is the cross-validated loss function, which is expected to take a U-shape owing to the bias–variance trade-off. We performed a grid search to find the optimal *h* that minimized the loss function.

### 2.3 Application to Breast Cancer Visium ST Data

To demonstrate the utility of our proposed method with real ST data, we analyzed a publicly available Visium ST dataset, downloaded from 10x Genomics (see the Data Availability Statement). The Visium Spatial Gene Expression Reagent Kit was used to generate the ST dataset from a section of breast cancer tissue taken from a woman in her 60s with a grade 3 tumor. The data included the horizontal and vertical coordinates of 2,518 detectable spots, and the median number of genes per spot was 5,244. We used CARD algorithm to obtain the cell-type proportions, and we used previously published scRNA-seq data (Wu et al. 2021) from patients with breast cancer as references. Nine cell types were defined in the reference dataset used in the CARD analysis: cancer epithelial cells, cancer-associated fibroblasts (CAFs), T cells, endothelial cells, perivascular-like cells (PVL), myeloid cells, B cells, normal epithelial cells, and plasmablasts. CAFs are highly specialized cells that communicate with cancer cells through the secretion of signaling molecules. Myeloid cells are differentiated descendants of hematopoietic stem cells in the bone marrow.

### 2.4 Simulation Study

We generated our simulation data to mimic the breast cancer ST dataset described in Section 2.3 in terms of spot location and cell-type proportions. The true CTS stemness of the *K* = 9 cell types, {*α*_*k*_(*u*_*i*_, *v*_*i*_); *k* = 1,2, … , *K, i* = 1,2, … , *N*} , were generated from bivariate Gaussian densities with various shapes: *α*_*k*_(*u*_*i*_, *v*_*i*_) was equal to the bivariate Gaussian density function evaluated at (*u*_*i*_, *v*_*i*_). The mean and variance of this density function took different values for different cell types so that the true CTS stemness could vary according to cell type and spatial location. Since stemness ranges from 0 to 1, we linearly transformed the Gaussian density to that range. The spot-level stemness was generated from the simulated CTS stemness, the cell-type proportions, a random residual error *ε* with a mean-zero uniform distribution, and a range that ensured the spot-level stemness was between 0 and 1. We simulated 100 datasets independently, estimated the *K* CTS stemness surfaces from each simulated dataset, and compared the average estimated CTS stemness with the true CTS stemness using correlation coefficients and scatter plots.

### 2.5 Shiny Application

Using R Shiny, we developed an interactive, user-friendly software application for computing and visualizing CTS stemness using ST data (**Figure 1A**). This online tool, which allows users to upload their own ST dataset and retrieve the output (cross-validation plots, stemness histograms, spatial plots of stemness, and numerical results), can be accessed through our GitHub site (see the abstract). The application features a statistical deconvolution model to calculate the CTS stemness value, a graphical user interface accompanied by a detailed tutorial with example data, and a series of graphical outputs. It does not require programming by the user, making it convenient for biological laboratories and medical researchers.

**Figure 1.**
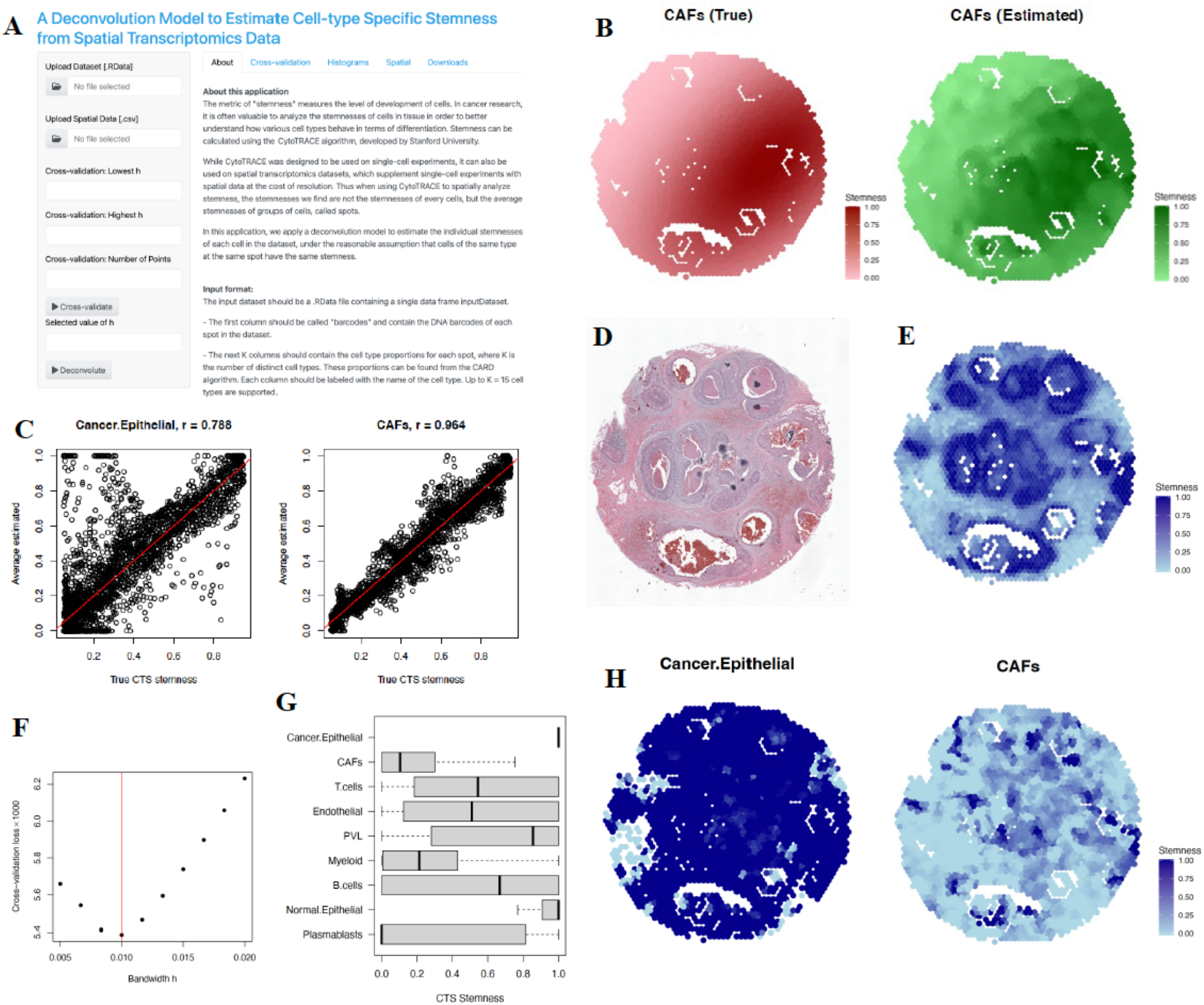
(A) The GitHub web application for the proposed method. (B) Spatial distribution of true vs. estimated cell type–specific (CTS) stemness of an arbitrarily selected cell type from the simulation study using the proposed method. (C) Correlation between the true and estimated CTS stemness of cancer epithelial cells and CAFs in the simulation. (D) The breast cancer tissue section from which the Visium spatial transcriptomic data were collected. The lumens are the interior spaces in white. (E) The spot-level stemness of the aforementioned tissue section. (F) Bandwidth vs. cross-validated loss function. The optimal bandwidth is indicated by the red line and corresponds to the minimized cross-validated loss function. (G) Distribution of estimated CTS stemness by cell type. (H) Spatial map of estimated CTS stemness of cancer epithelial cells and CAFs (spots with the corresponding cell type proportion < 0.1% excluded from the plot).

## 3 Results

### 3.1 Simulation Study

In the simulation study, we evaluated the accuracy of our method by comparing the estimated and the true CTS stemness. As we used the cell-type proportions from the real data, we named each cell type according to their actual names in the real data. The average true versus estimated CTS stemness surfaces for CAFs are shown in **Figure 1B**; those for the other cell types are shown in **Figure S1 A-C**. The areas of high estimated CTS stemness matched well with the areas of high true CTS stemness surfaces in all cell types. Scatterplots illustrating the correlation between the estimated and true CTS stemness for cancer epithelial cells and CAFs are shown in **Figure 1C**; those for the other cell types are presented in **Figure S2**. The correlation coefficient between the estimated and true CTS stemness across all cell types ranged from 0.788 (cancer epithelial cells) to 0.964 (CAFs) and was above 0.9 for 5 cell types. These results suggest that the proposed method can accurately deconvolve the CTS stemness from its spatial location.

### 3.2 Application to a Real Breast Cancer ST Dataset

Applying CytoTRACE to the spot-level gene expression Visium ST data generated from the breast cancer sample revealed that the pattern of spot-level stemness was similar to the darker areas in the histologic image (**Figure 1D** and **1E**), suggesting strong cell type– and tissue structure–associated stemness patterns. Most notably, cells surrounding lumens generally had higher stemness than those in other areas. The prevalence of different cell types in the tissue section varied substantially (**Figure S3**): CAFs, myeloid cells, normal epithelial cells, and cancer epithelial cells were among the most prevalent, whereas perivascular-like cells, B cells, endothelial cells, and plasmablasts were among the least prevalent. The proportion of each cell type also varied substantially across spatial locations (**Figure S4**).

To choose the optimal bandwidth for our local estimation, we plotted the relationship between the bandwidth and the cross-validated loss function (**Figure 1F**). We observed a U-shaped relationship between the bandwidth and the cross-validated loss function; based on that relationship, the optimal bandwidth was identified as *h* = 0.01. The distributions of estimated CTS stemness from all the spots by cell type are shown in **Figure 1G** and **Figure S5**. Epithelial cells had higher CTS stemness than other cell types. Cancer epithelial cells had higher CTS stemness than normal epithelial cells (two-sided *p*-value less than 0.001 from a Wilcoxon rank-sum test). In contrast, CAFs and myeloid cells had lower stemness than other cell types. T cells, B cells, plasmablasts, endothelial cells, and perivascular-like cells had stemness in between.

The estimated CTS stemness of cancer epithelial cells and CAFs varied with the spatial location (**Figure 1H**). (The estimated CTS stemness of the other cell types also varied with the spatial location [**Figure S6**].). Thus, one cell type can have substantial heterogeneity in stemness. A possible explanation for this finding is that the normal cell populations in our dataset have been impacted by the tumor cells and their transcriptomes were not completely normal. Many publications have reported strong field effects in tumor-neighboring areas, in which the normal cells possess characteristics and even genome aberrations that are similar to those of the tumor cells by which they are located (Martinez-Outschoorn et al. 2010, Chai and Brown 2009, Jones et al. 2005).

## 4 Discussion

We proposed a novel method for deconvolving CTS stemness from spot-level stemness. The method is flexible and has minimal modeling assumptions. Specifically, it allows the CTS stemness to vary by spatial location without any parametric distributions. Numerical studies supported the validity of the proposed method. As a general methodology, it can be used to deconvolve spot-level transcriptomics features other than stemness.

A potential limitation of the proposed method is that the deconvolution result depends on the estimated cell-type proportions. Systematic benchmark studies have shown that existing ST-based deconvolution methods, such as CARD and RCTD (Cable et al. 2022), achieve high accuracies if the ST and scRNA-seq references are matched (Li et al. 2022). However, if the matched scRNA-seq data are unavailable or of low quality, the accuracy of the estimated cell-type proportions may be affected and subsequently impact the performance of our method. Additionally, we took the estimated cell-type proportions as known values and ignored their potential variability. An approach that combines the cell-type deconvolution and CTS stemness analyses may achieve better accuracy and efficiency. Another limitation is that the tissue neighborhoods for the locally weighted analysis were selected on the basis of only mathematical considerations (i.e., the Euclidean distance between spots). A more refined approach would be to incorporate the tissue structure when defining the size and shape of local neighborhoods.

We believe that investigating the relationship between the estimated CTS stemness map and the tissue structure map is of scientific interest. For example, the stemness of a cell type may be affected by certain tissue features. The CTS stemness maps may also help in the investigation of cell-cell interactions, such as the relationship among cell groups with the same cell type but different stemness. Additionally, identifying stemness-associated CTS genes may also help provide a better understanding of CTS spatial heterogeneities. The proposed method serves as the first step toward these more extensive research directions.

## Funding

None

## Conflict of Interest

None

## Data Availability Statement

The real breast cancer Visium ST dataset analyzed in this paper is publicly available from 10x Genomics (https://www.10xgenomics.com/datasets/human-breast-cancer-ductal-carcinoma-in-situ-invasive-carcinoma-ffpe-1-standard-1-3-0).

## Notes

### Competing Interest Statement

The authors have declared no competing interest.

