## Supplementary document for "Delineating the Spatial Patterns of Cell Type–Specific Stemness Using a Statistical Deconvolution Model"

### Section S1. Stemness calculation using CytoTRACE

The metric of stemness, as calculated by CytoTRACE, is a ranking of all the spots in the datasets by differentiation status on a scale of 0 to 1, with 0 containing the least differentiation potential and 1 having the most differentiation potential. CytoTRACE analyzes the ST data as scRNA-seq data, ignoring the spatial information. The input to the algorithm is the genes expression counts at each spot. The algorithm identifies genes whose expression levels change in a consistent manner across spots; these genes are often associated with specific developmental stages or cell types. These genes are noted as possible predictors of cell development, and therefore differentiation. By comparing each spot's gene counts to the inferred trajectory, CytoTRACE can rank the spot based on relative levels of differentiation. Scaling these ranks down to a scale between 0 and 1 then gives the stemness of each spot.

### Section S2. Cell type proportions

Since the spot-level genetic feature is an ad-mixture of CTS genetic features, mixed by the proportions of cells in each type, it is necessary to know the cell type proportions in order to infer the CTS features (Rahmani et al 2019; Jin et al 2021; Feng et al 2022). We used the CARD algorithm (conditional autoregressive-based deconvolution; Ma and Zhou 2022), available in R package CARD, to calculate the cell type proportions in our dataset. Compared with previously proposed methods that were adapted from algorithms designed for scRNA-seq or bulk RNA-seq data (e.g., Andersson et al 2020; Cable et al 2022; Kleshchevnikov et al 2022), CARD makes use of the rich spatial localization information to identify these proportions: neighboring spots on the tissue likely contain more similar cell-type compositions than spots that are far away. CARD also uses the reference profiles of cell types, comparing the gene expressions in the dataset under study to the distinct genetic features of known cell types to determine the cell type composition.

**Section S3. Supplementary Figures**

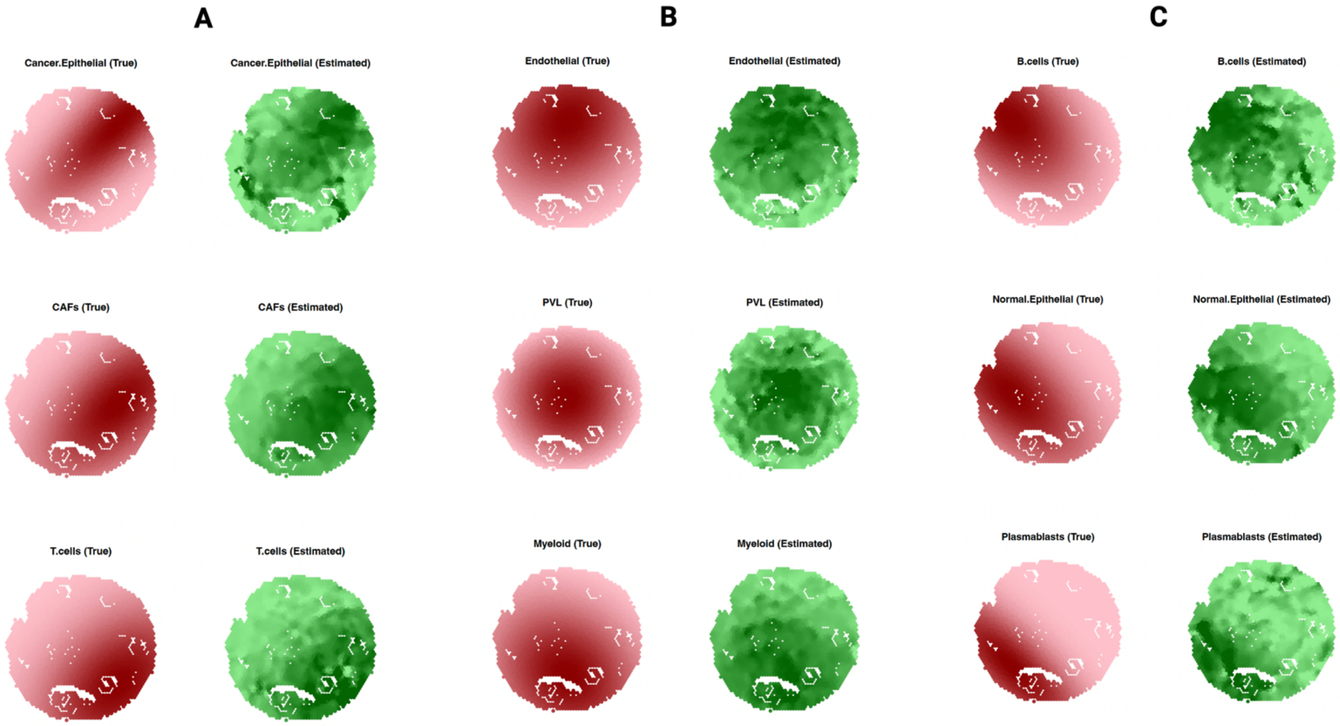


**Figure S1 (A)-(C).** The average estimated vs. the true CTS stemness surfaces for the 9 cell types in the simulation study. Plots (A)-(C) show three cell types each. The color scale is the same as **Figure 1B**.

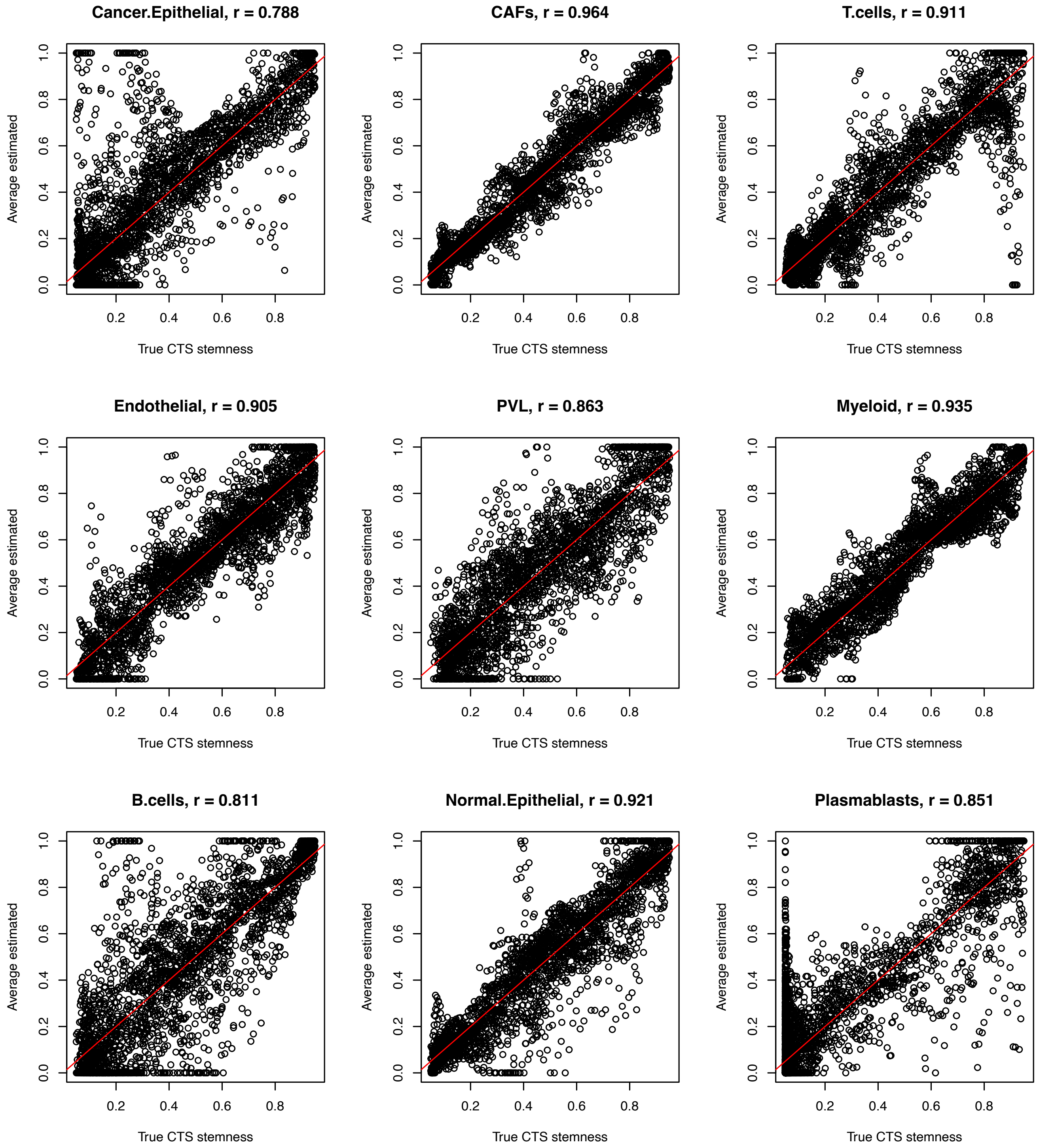


**Figure S2**. The average estimated vs. the true CTS stemness for the 9 cell types in the simulation study. The red line is the 45-degree line. The Pearson's correlation coefficient is also annotated with each plot.

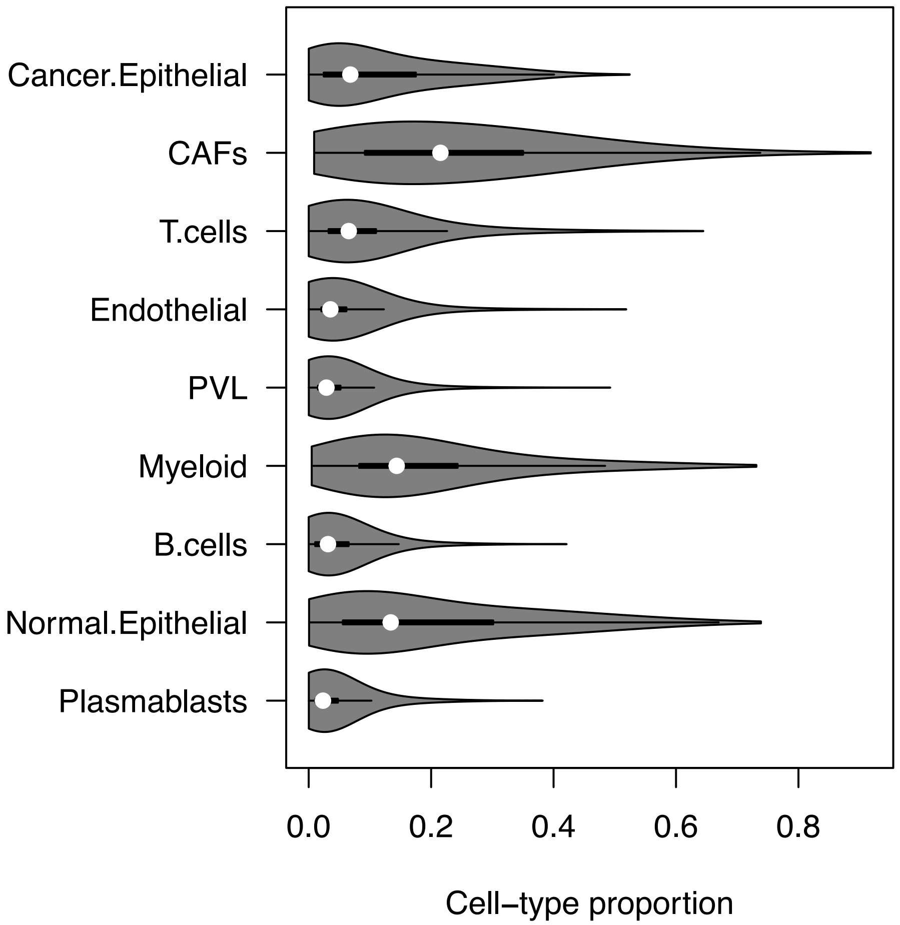


**Figure S3**. The distribution of the cell type proportions in a violin plot.

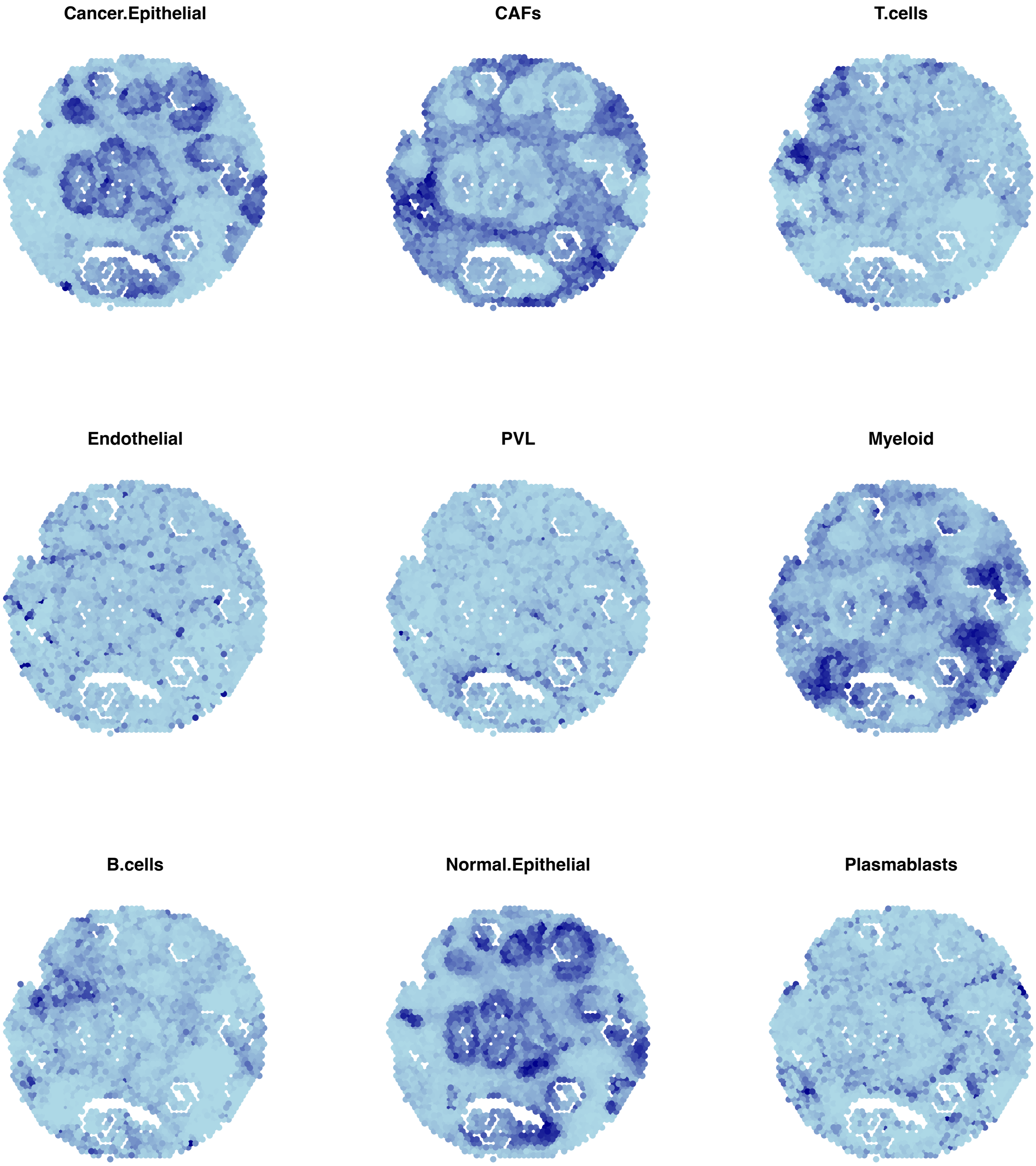


**Figure S4**. The spatial distribution of the cell type proportions by cell types and spot locations. The color scale is the same as **Figure 1H**.

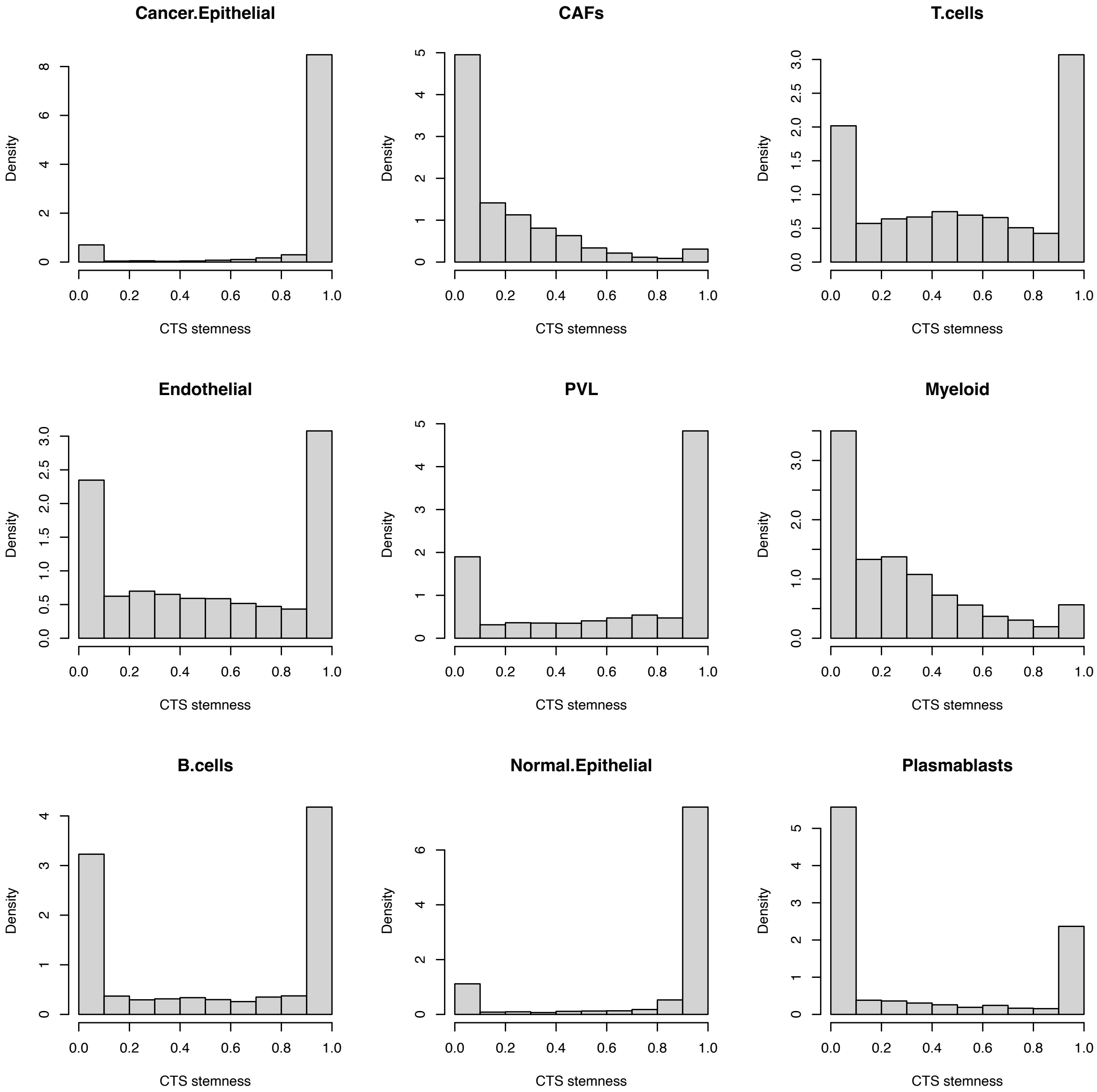


**Figure S5**. The histogram of the estimated CTS stemness by cell types.

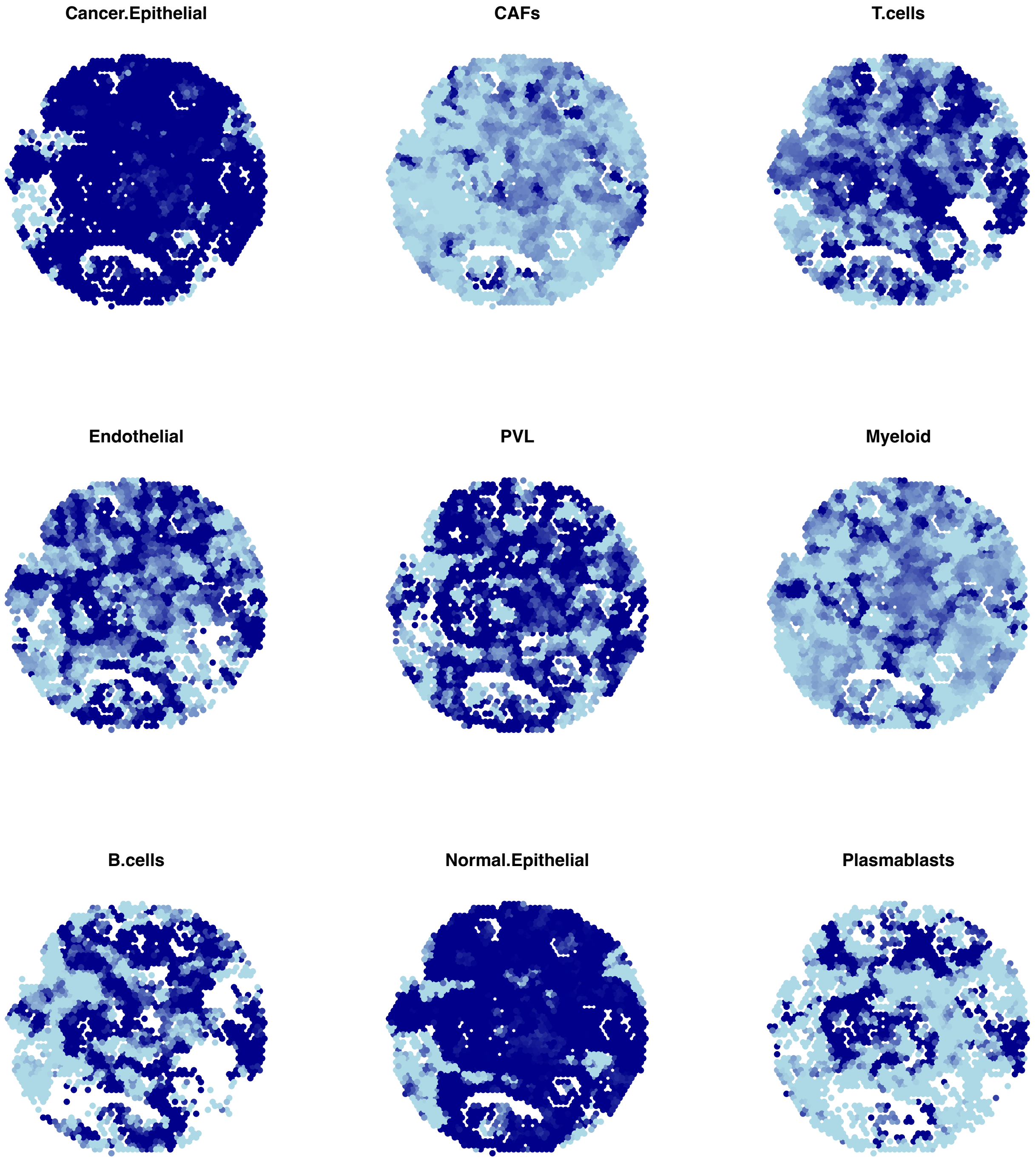


**Figure S6** The spatial distribution of the estimated CTS stemness by cell types and spot locations. The color scale is the same as **Figure 1H**.
